# Unraveling the Molecular Complexity of N-Terminus Huntingtin Oligomers: Insights into Polymorphic Structures

**DOI:** 10.1101/2024.02.12.579983

**Authors:** Neha Nanajkar, Abhilash Sahoo, Silvina Matysiak

## Abstract

Huntington’s disease (HD) is a fatal neurodegenerative disorder resulting from an abnormal expansion of polyglutamine (polyQ) repeats in the N terminus of the Huntingtin protein. When the polyQ tract surpasses 35 repeats, the mutated protein undergoes misfolding, culminating in the formation of intracellular aggregates. Research in mouse models suggests that HD pathogenesis involves the aggregation of N-terminal fragments of the Huntingtin protein (htt). These early oligomeric assemblies of htt, exhibiting diverse characteristics during aggregation, are implicated as potential toxic entities in HD. However, a consensus on their specific structures remains elusive.

Understanding the heterogeneous nature of htt oligomers provides crucial insights into disease mechanisms, emphasizing the need to identify various oligomeric conformations as potential therapeutic targets. Employing coarse-grained molecular dynamics, our study aims to elucidate the mechanisms governing the aggregation process and resultant aggregate architectures of htt. The polyQ tract within htt is flanked by two regions: an N-terminal domain (N17) and a short C-terminal proline-rich segment.

We conducted self-assembly simulations involving five distinct N17 + polyQ systems with polyQ lengths ranging from 7 to 45, utilizing the ProMPT force field. Prolongation of the polyQ domain correlates with an increase in *β*-sheet-rich structures. Longer polyQ lengths favor intra-molecular *β*-sheets over inter-molecular interactions due to the folding of the elongated polyQ domain into hairpin-rich conformations. Importantly, variations in polyQ length significantly influence resulting oligomeric structures. Shorter polyQ domains lead to N17 domain aggregation, forming a hydrophobic core, while longer polyQ lengths introduce a competition between N17 hydrophobic interactions and polyQ polar interactions, resulting in densely packed polyQ cores with outwardly distributed N17 domains. Additionally, at extended polyQ lengths, we observe distinct oligomeric conformations with varying degrees of N17 bundling. These findings can help explain the toxic gain-of function that htt with expanded polyQ acquires.

**Author summary:** Our study delves into Huntington’s disease (HD), a devastating neurodegenerative disorder triggered by abnormal expansions of polyglutamine repeats in the Huntingtin protein. When these repeats exceed a critical threshold, the protein misfolds, leading to the formation of harmful intracellular aggregates. Using computational techniques, we explored the intricate process by which these aggregates form and examined their complex structures.

Our findings shed light on the diverse nature of the protein fragments involved in HD pathology, emphasizing the importance of identifying various structural forms as potential targets for therapeutic intervention. We observed that changes in the length of the polyglutamine tract significantly impact the resulting aggregate structures, revealing insights into the disease mechanism. Specifically, we found that an expansion of the polyglutamine domain leads to distinct aggregate morphologies. In addition, the way the first 17 amino acids of these protein fragments pack against each other in the aggregates depends on the length of the polyglutamine repeats. By uncovering these structural intricacies, our study contributes to a deeper understanding of HD and may pave the way for the development of targeted treatments aimed at disrupting or preventing the formation of toxic protein aggregates.

## Introduction

The Huntingtin protein is a complex molecule with multiple domains, playing vital roles in various cellular processes such as axonal transport, signaling pathways, and protein interactions [1–3]. Notably, abnormalities in the glutamine repeat region (polyQ) located at the N terminus of the Huntingtin protein (htt) are associated with Huntington’s Disease (HD), a progressive neurodegenerative condition characterized by deteriorating physical and cognitive functions. In individuals without the disease, polyQ length ranges between 6 to 34 (htt^*wt*^), but clinical pathogenicity occurs when exceeding a threshold of 35Q residues (htt^*mutant*^). [4]. These elongated polyQ segments undergo misfolding, leading to the formation of intracellular aggregates, which are hallmark features of HD pathology. The length of the expanded polyQ domain correlates with earlier disease onset and increased disease severity, highlighting its significance in the progression of HD.

While Huntington’s Disease is among 9 recognized polyQ-based disorders, our understanding of the aggregation process and its underlying mechanisms remains incomplete [5]. PolyQ presents unique challenges for experimental study due to its rapid and length-dependent aggregation kinetics, conformational flexibility, and insolubility [6, 7]. Despite these difficulties, researchers have made progress in uncovering several structural characteristics of polyQ. PolyQ can self-assemble into polar zippers, where glutamine sidechains stabilize beta sheets [8]. Additionally, evidence suggests the presence of coiled-coil structures, *α*-helices, and *β*-helical conformations within the polyQ domain [9–11], illustrating its flexible nature. In solution, polyQ tends to be insoluble and exhibits a preference for self-interaction, resulting in collapsed and more globular structures as polyQ length increases [12, 13]. The structural and dynamic properties of polyQ are significantly influenced by its flanking sequences, particularly the N-terminal N17 domain and the C-terminal proline-rich domain (PRD) [14–16]. Studies indicate that N17 may adopt a coiled-coil state or an amphipathic helix, promoting htt aggregation [17, 18]. The helical nature of N17 can extend into the polyQ domain, initiating a region of helicity before polyQ adopts an extended structure [14].

Numerous aggregation models have been proposed, yet a consensus regarding the structure of aggregates remains elusive. Tryptophan fluorescence experiments on htt with varying polyQ lengths suggested that aggregates comprise an amyloid core involving both the polyQ and N17 domains [19]. Similarly, other studies propose ternary and quaternary interactions between N17 and polyQ at the fibril core, with the PRD extending from it [20, 21]. Conversely, ssNMR analysis of the N17Q_30_P_10_K_2_ fragment indicated that the amyloid core solely comprises the polyQ domain, while the N17 domain resides peripherally, more exposed to solvent [22]. Chen and Wolynes’ implicit solvent htt model for N17Q_20_ suggests N17 domain bundling, while the polyQ region forms trailing inter-peptide *β*-sheets [23]. Recent evidence suggests that smaller, polymorphic pre-fibrillar oligomers, rather than mature fibrils, are neurotoxic species in neurodegenerative disorders. However, characterizing them experimentally and structurally remains challenging due to their transient nature and the coexistence of monomeric, oligomeric, and fibrillar species.

A thorough comprehension of the aggregation process and oligomeric structures is imperative for the development of therapeutics targeting proteins. While techniques like cryo-EM and NMR have been instrumental in elucidating the morphology and polymorphisms of htt fibrils, molecular details of the oligomers remain elusive [20, 22, 24]. Molecular dynamics simulations offer a promising avenue to bridge these knowledge gaps, providing insights into the underlying molecular mechanisms.

Biological processes such as protein folding and aggregation occur over timescales inaccessible to unbiased all-atom molecular dynamics simulations. Hence, coarse-grained molecular dynamics, a reduced-resolution approach, becomes indispensable. By grouping local atoms into coarse-grained beads, this method yields smoother free energy landscapes, enabling simulations of larger systems over biologically relevant timescales.

In this study, we utilized our in-house developed ProMPT coarse-grained force field [25], featuring free-energy-based interaction parameterization and explicit dipoles to represent polar regions of the protein. Incorporating dipoles enables the force field to simulate protein structural transitions in response to the changes in local environment.

Our investigation focused on examining the monomeric and oligomeric states of both wild-type and mutant lengths of htt using coarse-grained molecular dynamics. We constructed five N17 + polyQ systems, comprising three wild-type (7Q, 15Q, and 35Q) and two mutant (40Q and 45Q) systems. Our results revealed a propensity for intra-molecular *β*-sheets as the polyQ length increased. We observed bundling of the N-terminal N17 domain in aggregate systems, which diminished with longer polyQ lengths. Furthermore, we characterized the influence of polyQ length on oligomeric morphology and the multiplicity of distinct oligomeric conformations. These variations in oligomeric morphology likely contribute to the observed polymorphism in fibrillar htt structures.

In this study, we have used coarse-grained molecular dynamics to examine the aggregation process and oligomer morphologies of both htt^*wt*^ and htt^*mutant*^. We observed bundling of the N17 domain in the aggregate systems, which reduces upon increase in polyQ length. At mutant polyQ lengths, there is a heightened variability in N17 bundling with N17 locating at the periphery of the aggregate. In contrast, N17 is located in the center of the oligomer for htt^*wt*^. In addition, distinct oligomeric conformations are observed for htt^*mutant*^ in contrast to htt^*wt*^. We suspect these variations in oligomeric morphology may be a structural basis for the polymorphism observed in fibrillar htt structures.

Our observations revealed the bundling of the N17 domain in aggregate systems, a phenomenon that diminishes with increasing polyQ length. Notably, at mutant polyQ lengths, we observed heightened variability in N17 bundling, with N17 predominantly located at the periphery of the aggregate. In contrast, in htt^*wt*^ systems, N17 tends to occupy the oligomer core. Additionally, we identified distinct oligomeric conformations in htt^*mutant*^ in contrast to htt^*wt*^. These variations in oligomeric morphology likely serve as a structural basis for the polymorphism observed in fibrillar htt structures.

## Materials and methods

All simulations were carried out on GROMACS 2019.4 simulator [26], visualised on VMD 1.9.4 [27], analyzed using in-house scripts using MDAnalysis version 1.1.1 [28]. Reweighting of simulation data was accomplished using the Multiscale Bennett Acceptance Ratio method, implemented in the pyMBAR package [29].

### Forcefield

We employed the Protein Model with Polarizability and Transferability (ProMPT) for all simulations, as detailed in our previous work [25]. ProMPT is a coarse-grained protein model designed to capture both secondary and tertiary structural changes. This capability is facilitated by the inclusion of dummy charges on all polar beads, allowing for the representation of structural transitions in response to local environmental changes. ProMPT has undergone validation on various small proteins exhibiting diverse structural characteristics [30]. Further information regarding parameterization and validation procedures can be found in the ProMPT paper.

For water representation, ProMPT was coupled with the MARTINI polarizable water model [31]. The coarse-grained representation of the N17 domain is illustrated in Fig. S1.

### Simulation Parameters

To characterize the aggregation process and oligomeric structure, six peptides were placed in a 12 nm box, initially in an unfolded state. In this model, a helical dihedral potential with competing minima was applied to the N17 backbone beads, utilizing a force constant of 10 kcal/mol (Fig. S2). Experimental evidence suggests that N17 adopts an unstructured or coil-like structure at low concentrations [19], while at high concentrations, it forms an amphipathic helix [32]. No dihedral potential was applied to the polyQ domain.

Restraints were applied to hold the peptides in place while the system underwent steepest descent energy minimization for 5000 steps, followed by a 5000 step NPT equilibration.Sodium and chloride ions, sourced from the MARTINI model, were introduced into the simulation box to achieve an effective salt concentration of 125 mM, mirroring typical experimental conditions for htt studies. After the addition of salt ions, a second round of energy minimization and equilibration was performed, consisting of 5000 steps each. Upon release of the constraints, a canonical production run extended for 350 nanoseconds, employing a timestep of 10 femtoseconds.

The peptide systems were subjected to a range of temperatures spanning from 380K to 500K, with increments of 20K, resulting in 7 temperatures for each peptide system. It’s important to note that the coarse-grained temperatures employed in this study are not directly comparable to real or experimental temperatures. However, varying the temperature enables a more thorough exploration of the free energy landscape, a common technique in molecular dynamics simulations [33–35]. Since these simulations were conducted under the NVT ensemble, the solvent density remained consistent across all temperatures. The Nose-Hoover thermostat, with a time constant of 1 picosecond, was utilized to maintain fixed temperatures. A relative dielectric constant of 2.5 was applied, and electrostatic calculations were performed using Particle Mesh Ewald. Constraints were evaluated using the LINCS algorithm, and the neighbor list was updated every 10 steps.

### Analysis Methods

To determine the probability of initial interactions between peptides, we focused on the first 90 nanoseconds of simulation time, corresponding to the period during which the oligomer formation occurred. Interactions between peptides were categorized as N17-Q, Q-Q, or N17-Q driven.

For subsequent analyses, we analyzed the last 260 nanoseconds of the simulations. This ensured that structural characterization was performed on segments of the trajectory where the complete oligomer had formed. Inter-peptide sheet contacts were quantified based on the number of contiguous contacts between the polyQ domains of different peptides, disregarding interactions within a single peptide. Intra-peptide sheet contacts referred to non-helical contiguous contacts within the polyQ domain of a single peptide. In both inter and intra-peptide sheet contacts, a minimum of 4 contiguous contacts was required to be classified as a sheet. The number of inter-peptide and intra-peptide contacts was then normalized by the maximum possible contacts in each system.

The hydrophobic solvent-accessible surface area (hSASA) was computed using the built-in *gmx sasa* tool from GROMACS. Specifically, hydrophobic residues F, L, M, and V were selected for analysis.

To determine the number of interacting N17 domains (*N*_*int*_), a distance matrix was constructed using the backbone and side chain beads of the N17 domain. Residues within 7 Å of each other were defined as being in contact. In order to filter out transient interactions of N17 domains, interactions were considered significant if there were at least 4 or more interactions between the domains.

To examine polymorphism in htt, trajectories were filtered based on the value of *N*_*int*_. Compactness was calculated as the ratio of the smallest and largest eigenvalues of the moment of inertia tensors, inspired by the implementation in ATRANET [36].

## Results

### Aggregation Pathway and Structural Characteristics

To elucidate the mechanisms driving aggregation, we assessed the likelihood of the initial interaction between the N17 and polyQ domains of the peptides (Fig. 1). Previous studies have suggested that the aggregation of polyQ is facilitated by the self-associating properties of N17. Consistently, in the 7Q system, initial inter-peptide interactions predominantly involve N17-N17 and N17-Q interactions. However, as the polyQ length increases, there is a notable increase in glutamine-driven interactions, such as N17-Q or Q-Q interactions, coupled with a decrease in N17-N17 driven associations.

**Fig 1.**
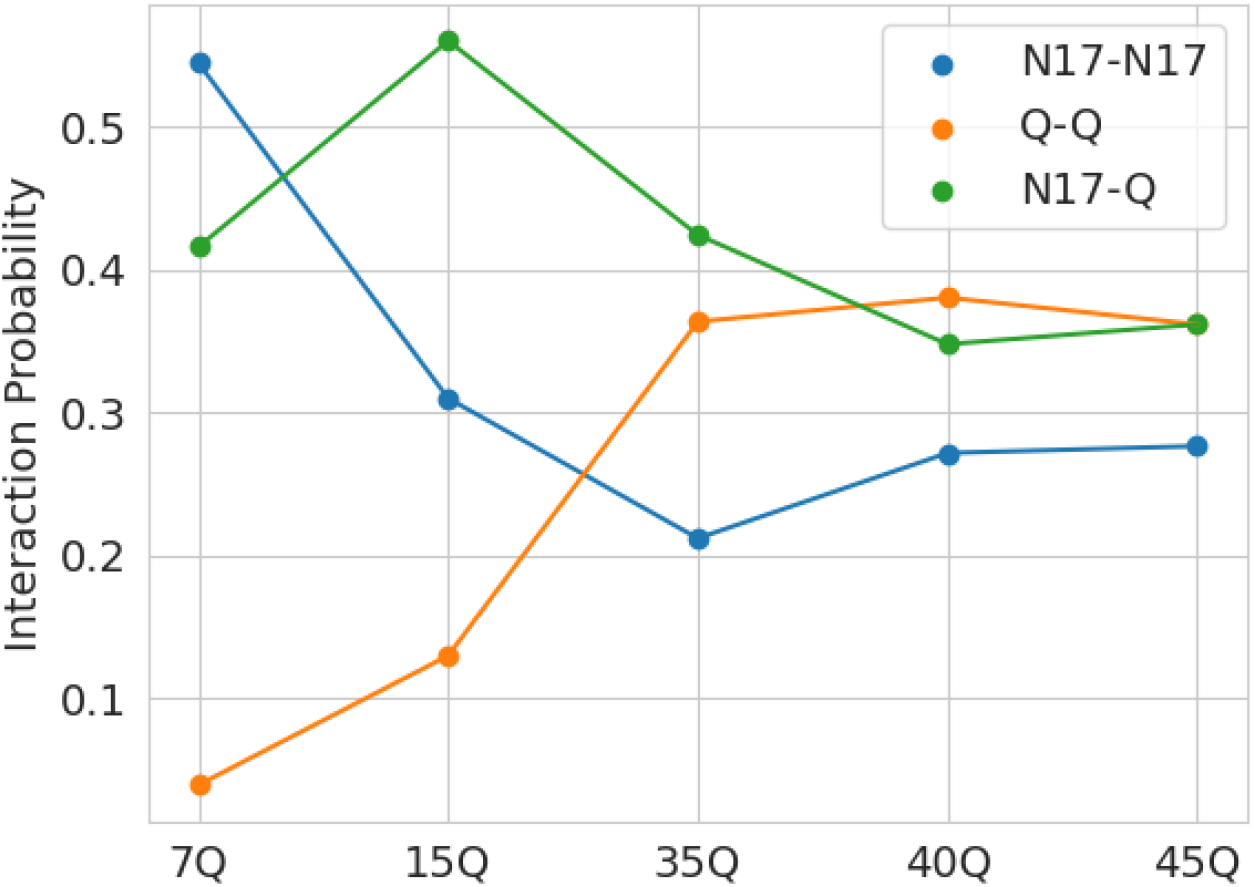
Probability of inter-peptide interaction occuring through N17-N17, N17-Q and Q-Q interactions.

Next, we delved into the structural characteristics of the peptide systems. Detailed information on monomeric structures is available in S1 File. Assessment of secondary structure characteristics in the oligomers was conducted by examining helical fraction and *β*-sheet fraction. We observed that the N17 domain predominantly adopts helical conformations, with a helical fraction of 0.9, while the polyQ domain exhibits a helical fraction of approximately 0.2 across all polyQ lengths, as depicted in Fig. S3. In terms of *β*-sheet formation (Fig. 2), minimal to no *β*-sheet formation was observed in the 7Q system due to the short length of the polyQ domain. In the 15Q system, inter-peptide *β*-sheets were favored over intra-peptide *β*-sheets. A snapshot of the 15Q system (Fig. 2C) illustrates the alignment of polyQ domains between peptides, forming inter-peptide *β*-sheets, albeit with less frequent intra-peptide *β*-sheet formation. However, as the polyQ length increases (polyQ ≥ 35), the preference shifts towards intra-peptide *β*-sheets, as demonstrated in a snapshot of the 45Q system (Fig. 2D).

**Fig 2.**
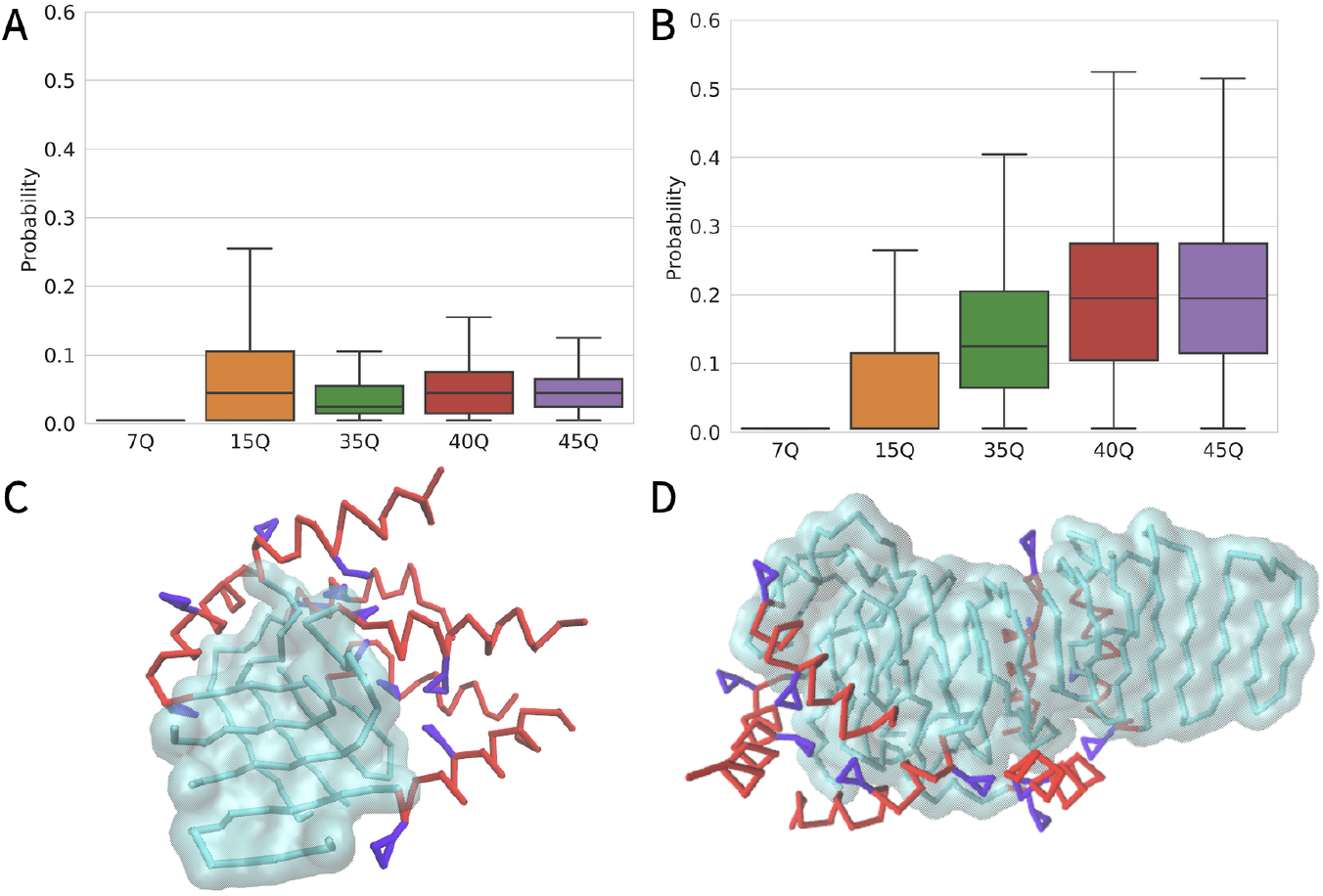
Beta sheet preferences of the polyQ domain, reweighted to 380K. A) Normalized inter-peptide *β*-sheet contacts, B) Normalized intra-peptide *β*-sheet contacts, C) Snapshot of inter-peptide *β*-sheets observed in the 15Q system, D) Snapshot of intra-peptide *β*-sheets observed in the 45Q system. N17, polyQ domain and F side chain residues are colored in red, cyan and violet respectively.

These observations are consistent with the findings of Yushchenko *et al*. [37], who utilized FTIR spectroscopy to analyze htt fragments with different polyQ lengths, and with the research by Bugg *et al*. [20], which emphasized the prevalence of anti-parallel *β*-sheets. Moreover, various biophysical investigations have highlighted the polyQ domain’s conformational flexibility, demonstrating its ability to adopt helices, coiled coils, and *β*-sheet-rich structures [9–11, 38].

It is noteworthy that our simulations did not indicate the presence of coiled coils, contrary to the findings of Chen and Wolynes [23], who observed extensive inter-peptide *β*-sheets in simulations involving N17Q_20_ and identified coiled-coil structures in the N17 domain. The snapshots provided in Fig. 2 illustrate that larger polyQ domains tend to collapse upon themselves, resulting in an increase in intra-peptide *β*-sheets and the formation of hairpin-like structures. This preference for more collapsed configurations is evident not only in the oligomeric systems but also in the monomeric systems (see S1 File). Several studies have indicated that longer polyQ domains tend to adopt more collapsed, globular structures [13, 39]. Recently, Kang *et al*. demonstrated this phenomenon with varying lengths of polyQ in htt [13].

### Competition between Hydrophobic and Polar Interactions

Upon visual examination of the oligomers, we observed the presence of a hydrophobic core in systems with shorter polyQ lengths. However, as the polyQ length increased, the aggregation of hydrophobic interactions diminished, resulting in the loss of the hydrophobic core. To quantify this trend, we plotted the average distance of phenylalanine (F) beads from the oligomer’s center of mass (Fig. 3A). As anticipated, the distance between F residues and the oligomer’s center of mass increased with longer polyQ lengths, indicating that F and other hydrophobic residues in the N17 domain were pushed towards the periphery of the oligomer, while the core comprised predominantly the polyQ domain.

**Fig 3.**
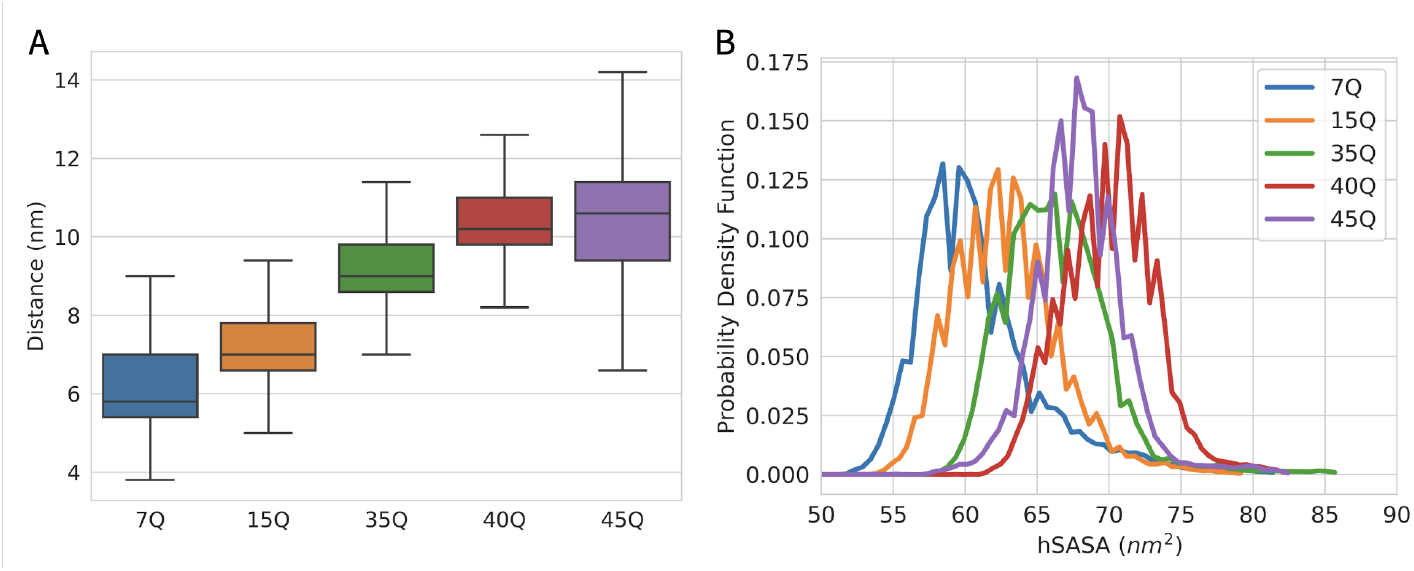
A gradual loss of the hydrophobic core with increasing polyQ length occurs due to the re-positioning of N17 to the periphery of the oligomer. A) Distance between the center of of mass of F sidechain beads from the center of mass of the whole aggregate. B) Hydrophobic Surface Accessible Solvent Area of the oligomers of all peptide systems.

Furthermore, to assess the environment surrounding the N17 domain, we examined the Hydrophobic Solvent Accessible Surface Area (hSASA) (Fig. 3B). A gradual increase in hSASA was observed with longer polyQ lengths, suggesting that as the polyQ length increased, the hydrophobic residues in the N17 domain became progressively more exposed to solvent.

The repositioning of N17 to the periphery of the aggregate can be attributed to a competition between hydrophobic interactions within the N17 domain and polar interactions involving the Q residues. In systems with shorter polyQ lengths, the dominance of hydrophobic interactions among N17 domains leads to their bundling. However, as the polyQ length increases, the influence of polar Q-Q interactions becomes more prominent, diminishing the bundling of N17 domains due to a reduction in the dominance of hydrophobic interactions.

The same trend is evident in the radius of gyration of the phenylalanine (F) sidechains, as depicted in Fig. S5, which increases with polyQ length. At shorter polyQ lengths, the hydrophobic residues tend to face inward due to N17 bundling, resulting in lower values of the radius of gyration. Conversely, in systems with longer polyQ domains and consequently reduced N17 bundling, the F sidechains become more exposed to solvent, leading to higher values of the radius of gyration. Short videos depicting the final oligomeric structures for polyQ lengths of 15 and 45 are provided in Video S1 and Video S2, respectively. Previous studies on the N17Q_30_P_10_K_2_ fragment of htt using solid-state nuclear magnetic resonance (ssNMR) techniques have suggested an external distribution of N17 [22].

### Variations in Oligomer Morphology and Observed Polymorphisms

To gain insight into the possible conformations of oligomeric structures, we evaluated the number of interacting N17 domains within the oligomer, denoted as *N*_*int*_. With 6 peptides in our simulation, *N*_*int*_ can range from 0 to 6, except for 1. Fig. 4A illustrates the probability distribution of various *N*_*int*_ values for each peptide system. At shorter polyQ lengths, there is a high probability of observing *N*_*int*_ = 6, indicating extensive bundling between N17 domains within the oligomer. For instance, in the 7Q system, the probability of *N*_*int*_ = 6 is approximately 0.55. However, as the polyQ length increases, this probability decreases, with the 45Q system showing the highest probability for *N*_*int*_ = 2.

**Fig 4.**
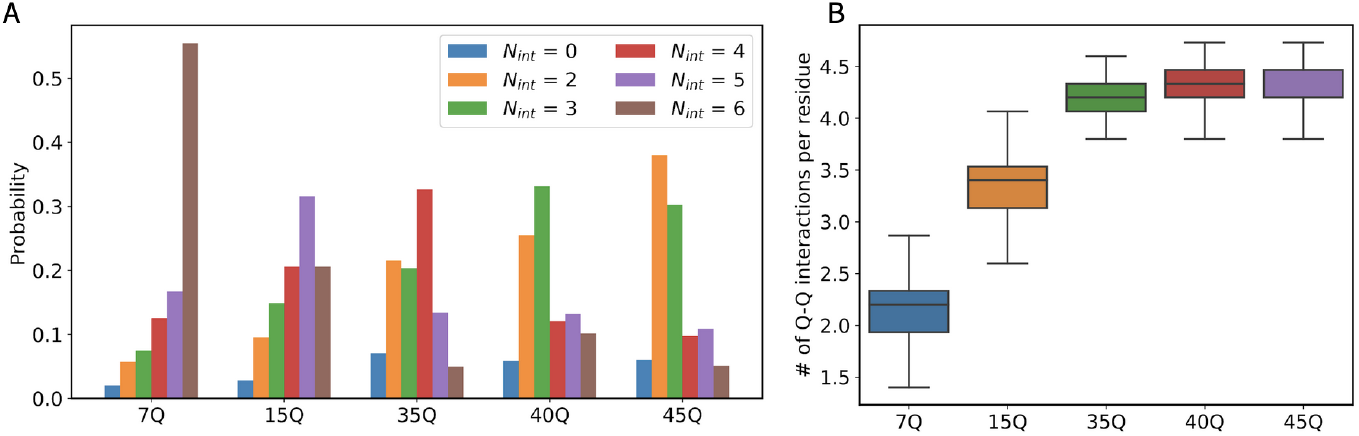
Oligomer morphology depends on polyQ length. A) Probabilities of the number of interacting N17 domains (N_*int*_) in each protein system. B) Number of Q-Q interactions per residue for each peptide system.

For htt^*mutant*^, we observe a heightened variability in the number of interacting N17 domains compared to htt^*wt*^, indicating multiple oligomeric conformations with distinct N17 distributions on the oligomeric surface. Previous experimental studies have demonstrated that N17 engages in tertiary and quaternary interactions involving both hydrophobic and hydrophilic interfaces at longer polyQ lengths [20]. This aligns with the nonspecific bundling of N17 observed in our simulations, reflected in the high variability of *N*_*int*_ and solvation of F sidechains. Despite its amphipathic nature driving N17 to bundle with neighboring domains through hydrophobic interactions, this tendency is hindered by increasing polyQ length.

Recent cryo-electron microscopy (cryo-EM) studies by Nazarov *et al*. have suggested that the core of htt fibrils consists of heterogeneous structures within the polyQ core, while the N17 and proline-rich domain (PRD) remain highly mobile around the polyQ core. Although concrete conclusions await a 3D cryo-EM structure, the reduction in *N*_*int*_ observed in our simulations with long polyQ aligns strongly with their findings [24].

In tandem with the fluctuations in *N*_*int*_, there is an augmentation in the average number of Q-Q interactions per glutamine residue, as seen in Fig. 4B. In polyQ systems with Q≥ 35 each glutamine residue interacts, on average, with 4 or more other glutamine residues. This is indicative of a vast polar, dipole bonding network between the glutamine residues, consistent with the polyQ domain’s reported capacity to form hydrogen bonds (which are a kind of dipole interaction) via its side chains and its propensity for self-association [8]. These trends align with the results of the monomeric simulations, where the average number of Q-Q interactions per glutamine residue increases, while the average number of water beads interacting with each glutamine residue reduces (see S1 File). These findings are in agreement with simulations of polyQ performed by Wen *et al*. and a growing body of work that posit water is a poor solvent for polyQ [40–42]

To further elucidate the diverse oligomeric conformations characterized by distinct N17 locations, we analyze the shapes of the oligomers, as illustrated in Figure 5. A compactness value of 1 indicates a spherical aggregate, while a value closer to 0 represents a prolate/extended conformation.

**Fig 5.**
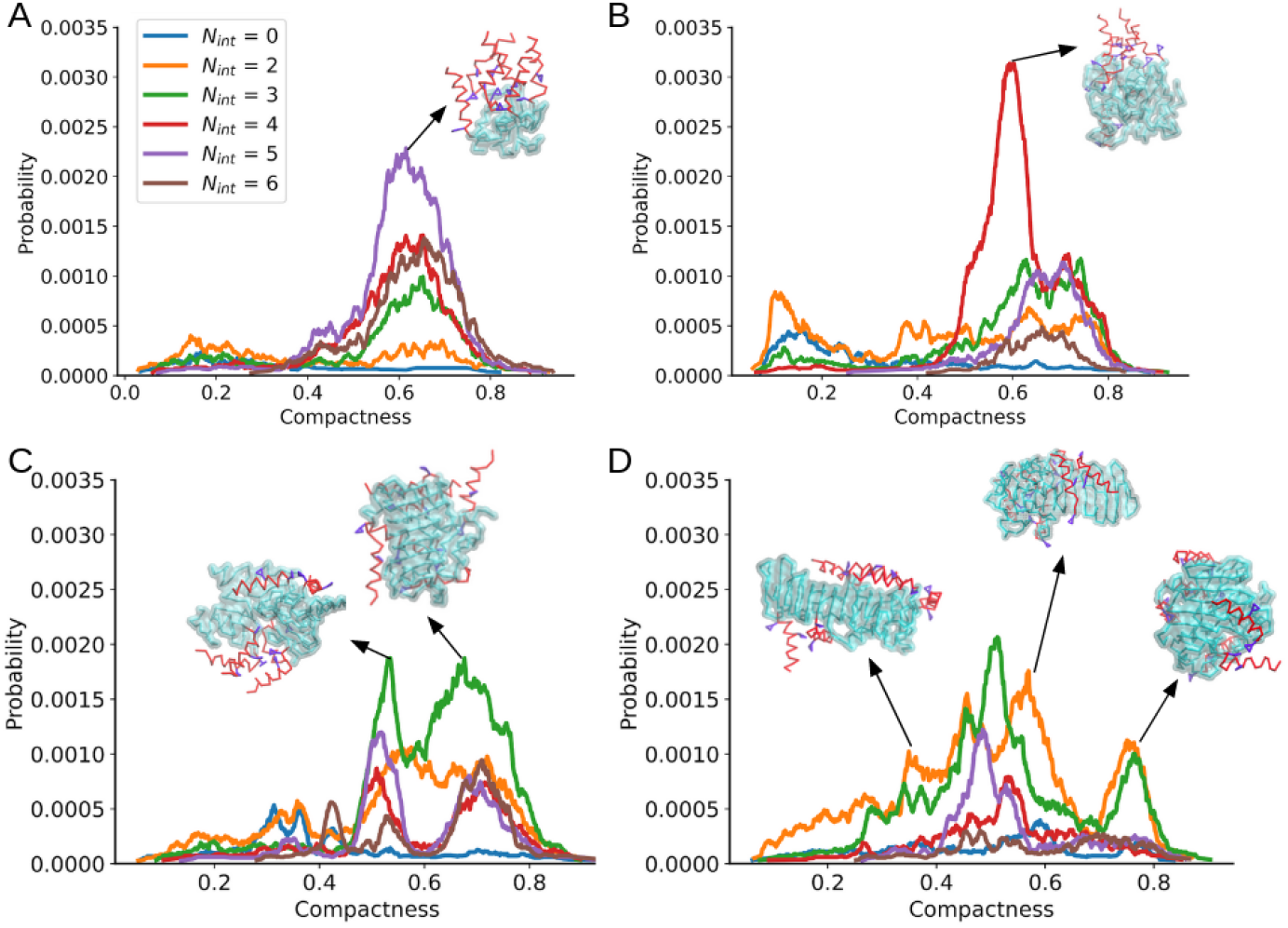
Flexibility in oligomer shape is indicative of polymorphism observed in systems with long polyQ lengths. Oligomer compactness distributions for A) 15Q, B) 35Q, C) 40Q and D) 45Q systems at different values of the number of interacting N17 domains (N_*int*_). N17, polyQ and F side chains are represented in red, cyan and violet respectively.

The compactness of oligomers reveals a notable correlation with the length of the polyQ segment. In the 15Q system (Figure 5A), a singular peak around a compactness value of approximately 0.58 is consistently observed across all *N*_*int*_ values, indicating a predominant polyQ oligomeric conformation unaffected by the number of interacting N17 domains. The inset snapshot illustrates a representative image for the 15Q system with *N*_*int*_ = 5, showing bundlings of the N17 domain resulting in a hydrophobic core, evident from the close proximity of F side chains (violet). The 7Q system demonstrates a similar trend (see Figure Fig. S5).

As the polyQ length increases, the oligomer can adopt more extended conformations, as evidenced by the broadening of the compactness distributions. The 40Q system presents a bimodal distribution for *N*_*int*_ > 2, suggesting two major conformations adopted by the polyQ domain in the oligomeric structure. Representative snapshots corresponding to the peaks of *N*_*int*_ = 3 are presented. In the 45Q system, there is an overall increase in the variety of shapes the oligomer can take on, as evident from the multiple peaks in the compactness distribution. The snapshots presented in Figure 5D correspond to regions of *N*_*int*_ = 2. The oligomer can adopt a range of conformations, from spherical to elongated shapes. The ensembles with low values of *N*_*int*_ (*N*_*int*_ < 4) have broader distributions, adopting conformations that can be either spherical or elongated. An enhanced variability in the shape of the oligomer is observed because low *N*_*int*_ values are observed at longer lengths of polyQ (Figure 4 A).

The polymorphic nature of htt emerges predominantly at longer polyQ lengths, enabling the oligomer to adopt multiple conformational states corresponding to varying levels of compactness and distinct positioning of N17 domains. Conversely, in oligomers with shorter polyQ lengths, characterized by primarily high *N*_*int*_ values, the range of conformational polymorphisms is considerably restricted, resulting in overlapping compactness distributions.

These findings underscore the conformational flexibility and polymorphic nature of htt protein fragments. At shorter polyQ lengths, the oligomers adopt more uniform structures, as evidenced by the probability of having high *N*_*int*_ (i.e., finding all N17 domains interacting) as seen in Figure 4, and the low variability in compactness (Figure 5). Here, the hydrophobic interactions between N17 domains lead to a more invariable oligomer. In contrast, systems with longer polyQ lengths experience two contrasting forces – hydrophobic interactions between N17 and the polar interactions between the Q residues. These opposing interactions result in oligomers that can adopt a wider variety of conformations, as seen by the numerous peaks in the compactness distributions. So far, the current knowledge on polymorphism has come from studying fibrillar structures. We hypothesize that these results may provide insight into the molecular mechanisms behind the varying amyloid states.

## Discussion

We observe a shift in the driving forces of association as polyQ length varies: shorter polyQ lengths predominantly exhibit N17-N17 interactions, whereas longer polyQ lengths show an increased prevalence of Q-Q/N17-Q interactions between peptides. Our findings align well with existing experimental studies, where the polyQ domain demonstrates a propensity to form helices and sheets. Moreover, the length of the polyQ sequence dictates the preference for inter-molecular sheet contacts over intra-molecular ones, particularly favored at longer polyQ lengths. These results corroborate spectroscopic findings by Yushchenko *et al*. [37].

We have revealed how oligomeric morphology varies with the length of the polyQ segment. Short polyQ lengths lead to oligomers with a distinct hydrophobic core, driven by the bundling of N17 domains where their hydrophobic sides face inward. As the polyQ segment elongates, N17 bundling diminishes, and the oligomer core transitions to interlinked glutamine regions, while N17 relocates to the aggregate surface. This finding aligns with previous studies indicating tertiary and quaternary interactions among N17 domains [20] and N17 positioning on the periphery of the amyloid core [22]. The repositioning of N17 results from a competition between its hydrophobic effects and the polar interactions of the polyQ tract, with hydrophobic interactions dominating at shorter lengths. We speculate that these insights could shed light on the heightened toxicity observed in htt^*mutant*^ aggregates.

A number of studies have indicated that in Huntington’s disease, the protein undergoes a toxic gain of function by interfering with normal function. One such mechanism involves the sequestration of transcription factors like CREB binding protein by mutant htt [43]. Co-aggregation of htt with CREB, increased cell toxicity, and depletion of CREB binding protein have been reported [44, 45]. Notably, our findings reveal that in htt^*mutant*^ oligomers, N17 is significantly more solvent exposed (see Fig. 3). This heightened exposure, particularly of the hydrophobic F residues in the N17 domain, could create favorable binding sites. We hypothesize that these oligomers may bind to essential enzymes and proteins, leading to the depletion of cellular resources crucial for maintaining homeostasis.

Furthermore, our study demonstrates that htt^*mutant*^ oligomeric structures exhibit polymorphic characteristics. A measure of oligomer shape was employed to compare ensembles with similar *N*_*int*_ values (representing the number of interacting N17 domains) within peptide systems. For systems with longer polyQ (≥35), multiple shapes are observed from spherical to elongated. In contrast, for polyQ < 35 predominantly displayed overlapping compactness distributions for all *N*_*int*_ values, indicating that htt^*wt*^ oligomers mainly adopt a single conformation that remains consistent across changes in *N*_*int*_.

Amyloid fibrils are known to exhibit various conformations from the same protein sequence, allowing them to propagate through the recruitment of additional molecules [46, 47]. This process typically involves the exposure of hydrophobic residues to the solvent. In on-pathway folding, these hydrophobic interactions stabilize the protein core, whereas in off-pathway folding, they can recruit other proteins [48, 49]. We hypothesize that the reduction in N17 bundling may contribute to the polymorphisms observed in htt^*mutant*^ fibrils.

Moreover, N17 has been identified as one of the membrane-binding domains of htt, facilitating its subcellular localization to various organelles [50]. It contains a nuclear export sequence that directs its translocation into the cytoplasm [51]. Chaibva *et al*. discovered N17’s ability to sense membrane curvature [52]. Therefore, the external distribution of N17 in htt^*mutant*^ oligomers could explain a potential gain-of-function mechanism in diseased states, particularly its interaction with membranes. Our future research will involve characterizing htt^*wt*^ and htt^*mutant*^ oligomers in association with lipid membranes of varying curvatures and compositions. Additionally, we will investigate the effects of the C-terminal proline segment of htt on aggregate morphology.

## Supporting information

**S1 File. Monomeric htt simulations**. Representative images, asphericity calculations, contact maps, domain specific interactions.

**Fig. S1.**
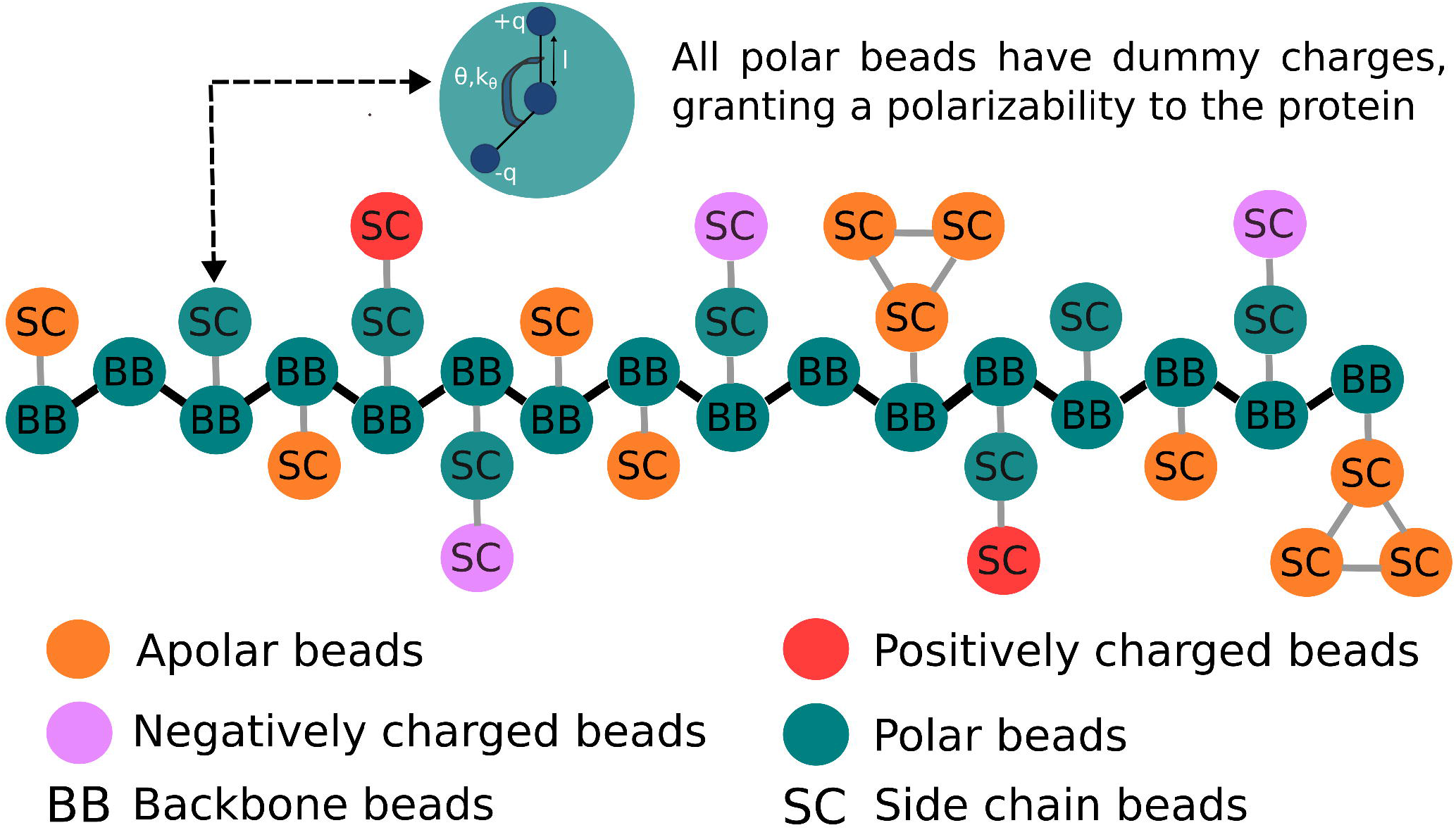
Coarse grained representation of the N17 domain. The forcefield was implemented as described in the ProMPT paper [25], with the exception of cation pi interactions.

**Fig. S2.**
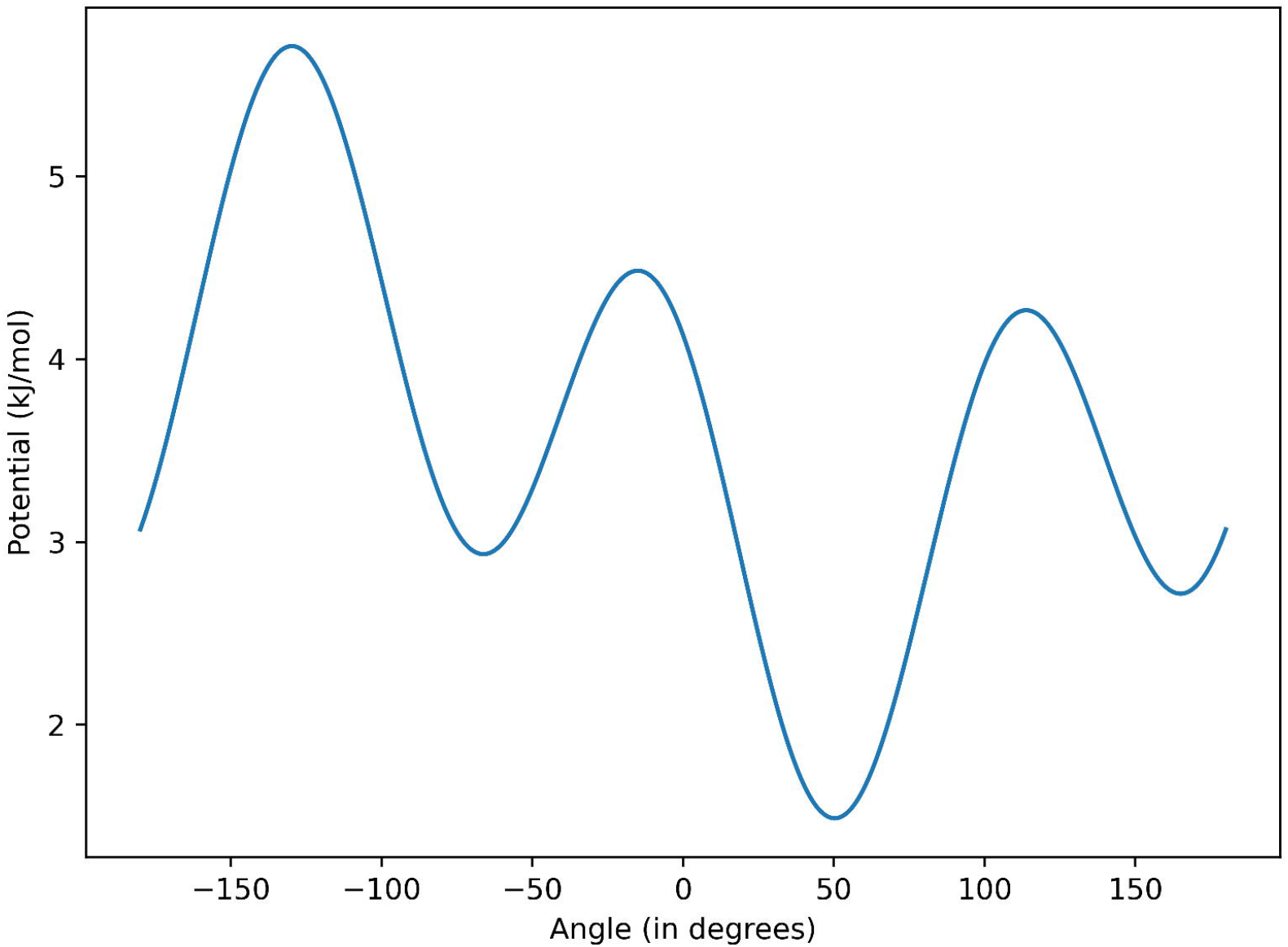
Dihedral potential applied to the backbone beads of the N17 domain.

**Fig. S3.**
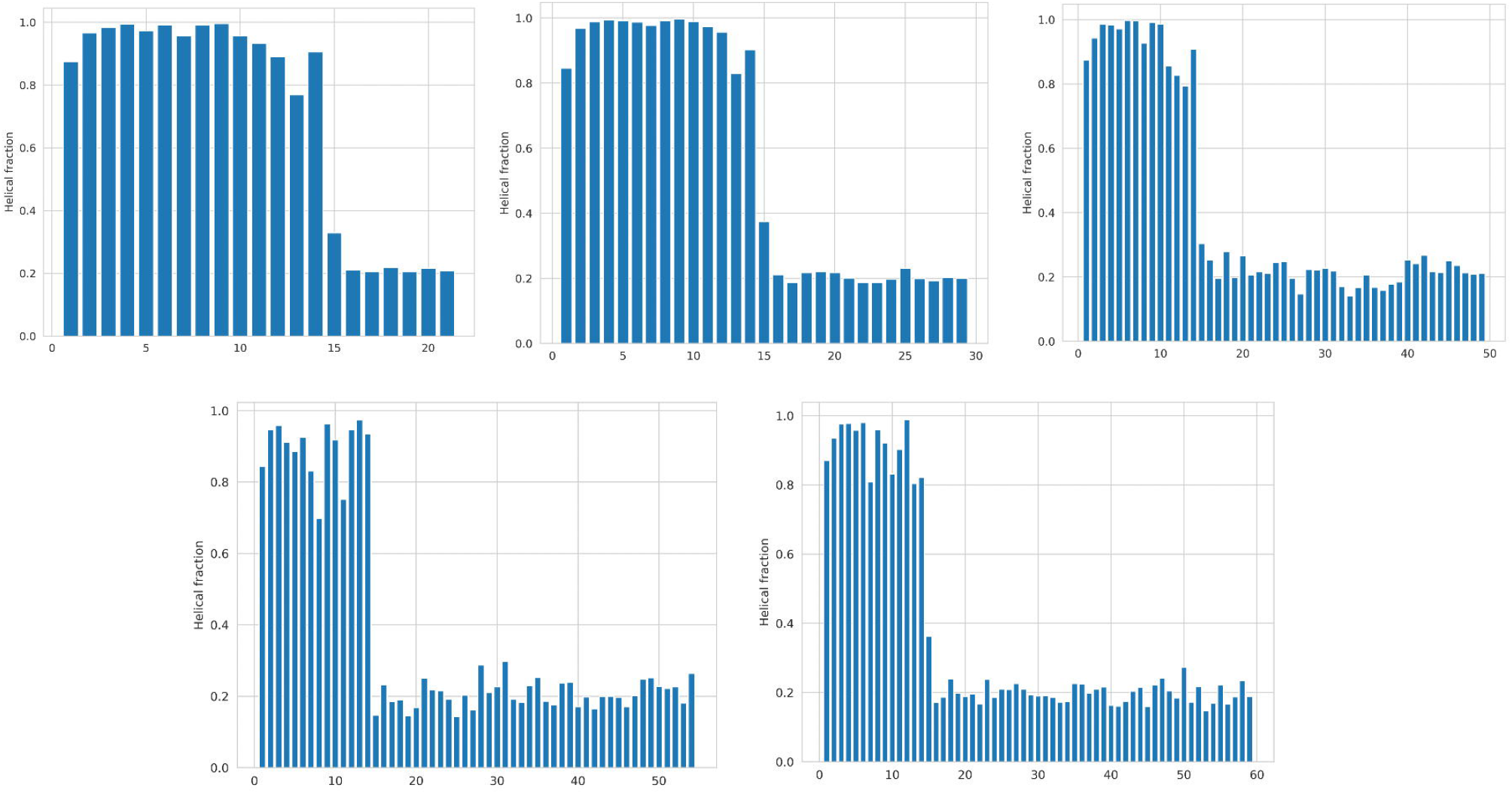
Helical fraction for oligomeric peptide systems. A) N17+7Q, B) N17+15Q, C) N17+35Q, D) N17+40Q, E) N17+45Q.

**Fig. S4.**
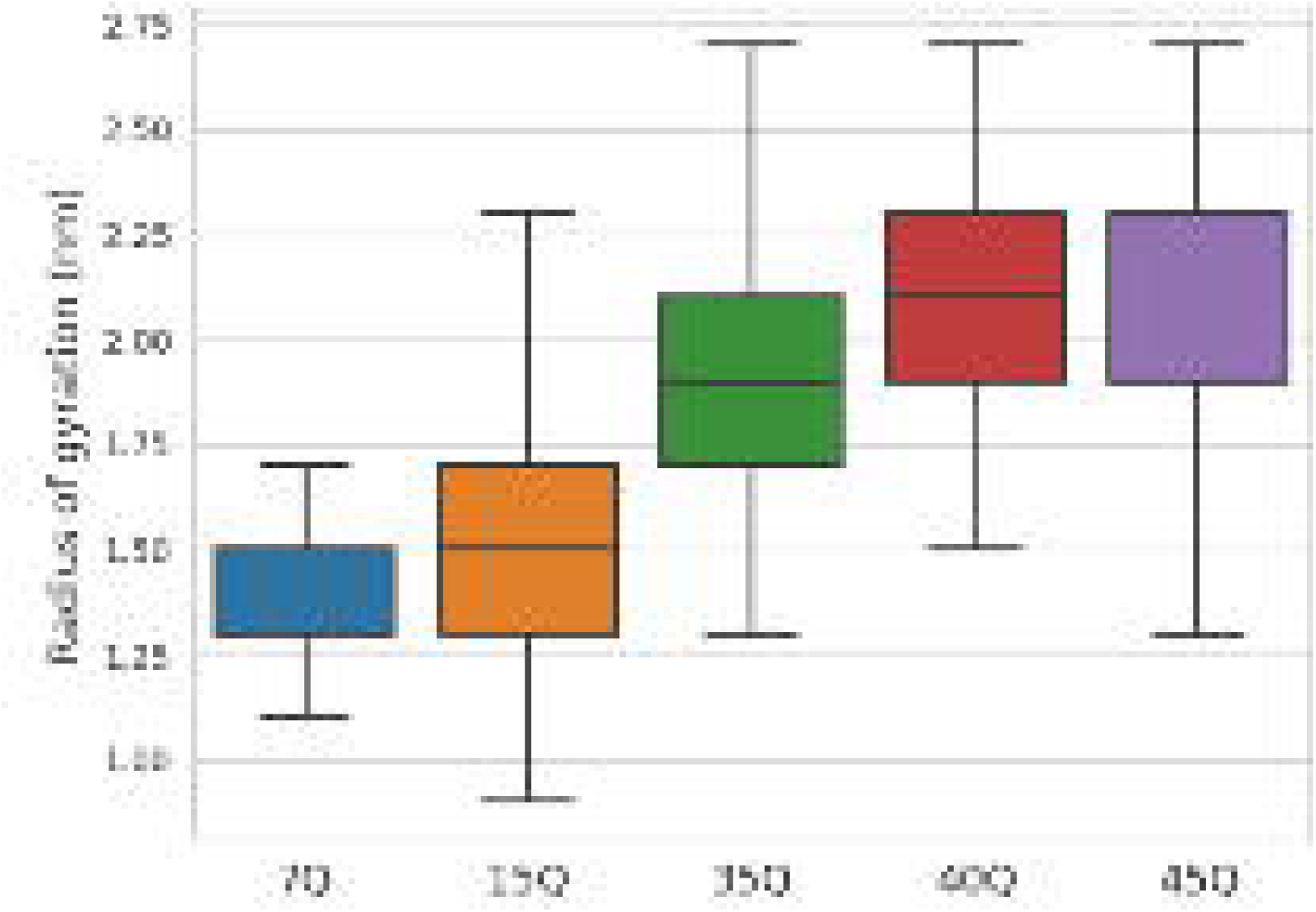
Radius of gyration of the F side chain beads.

**Fig. S5.**
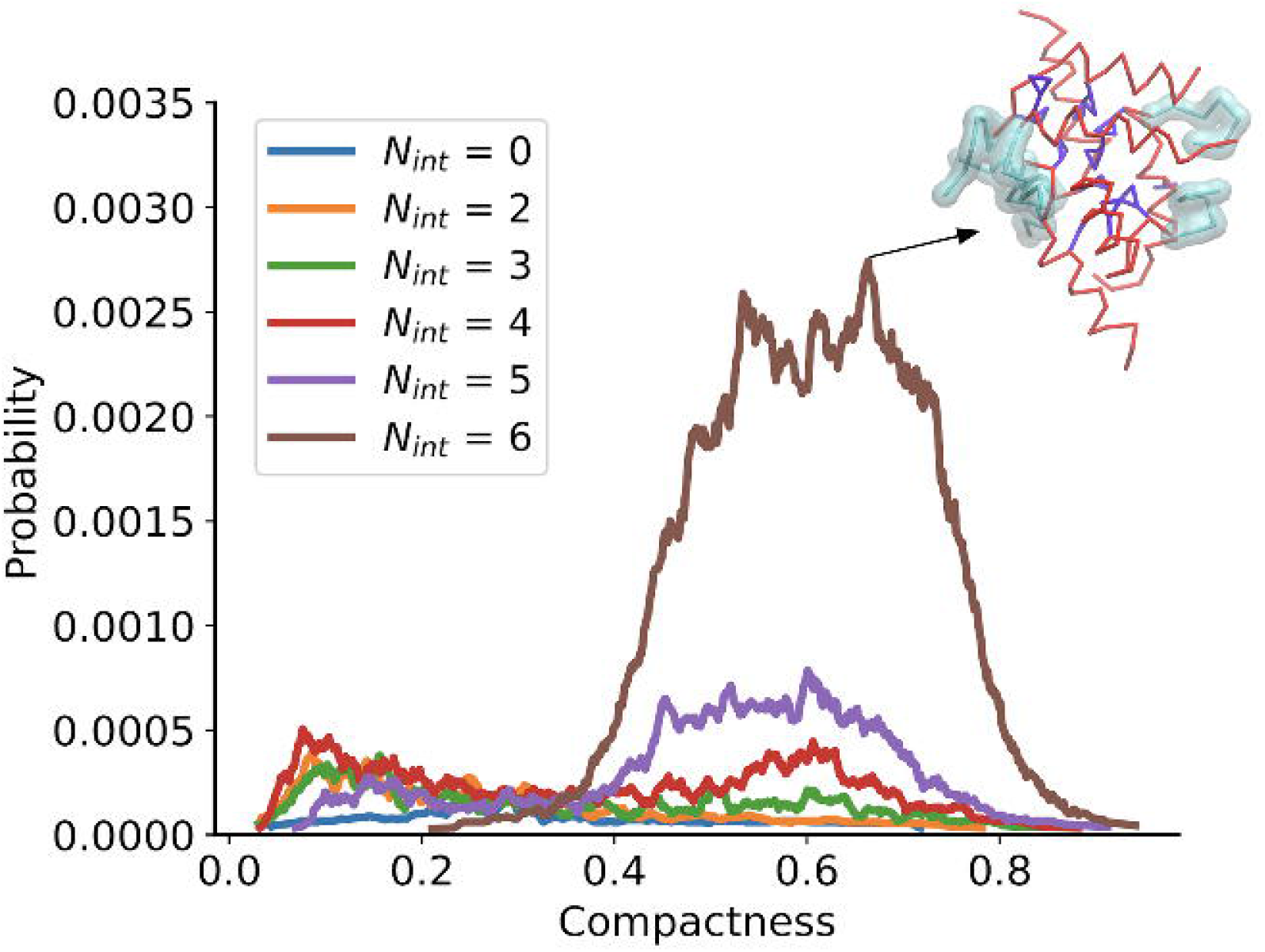
Compactness analysis for 7Q system.

**Video S1.**
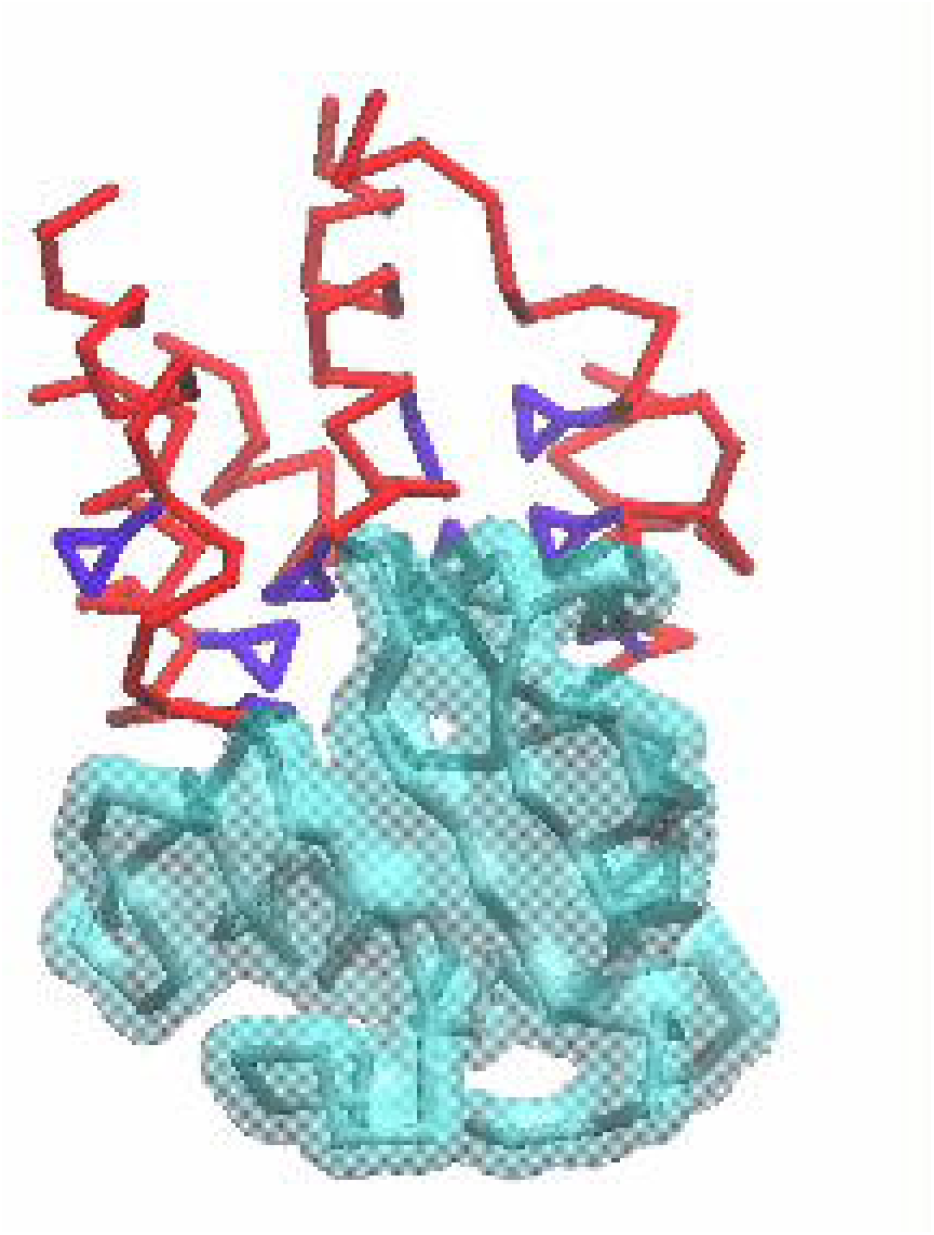
Bundling of N17 domains in the 15Q system. N17, polyQ and F are denoted in red, cyan and violet respectively.

**Video S2.**
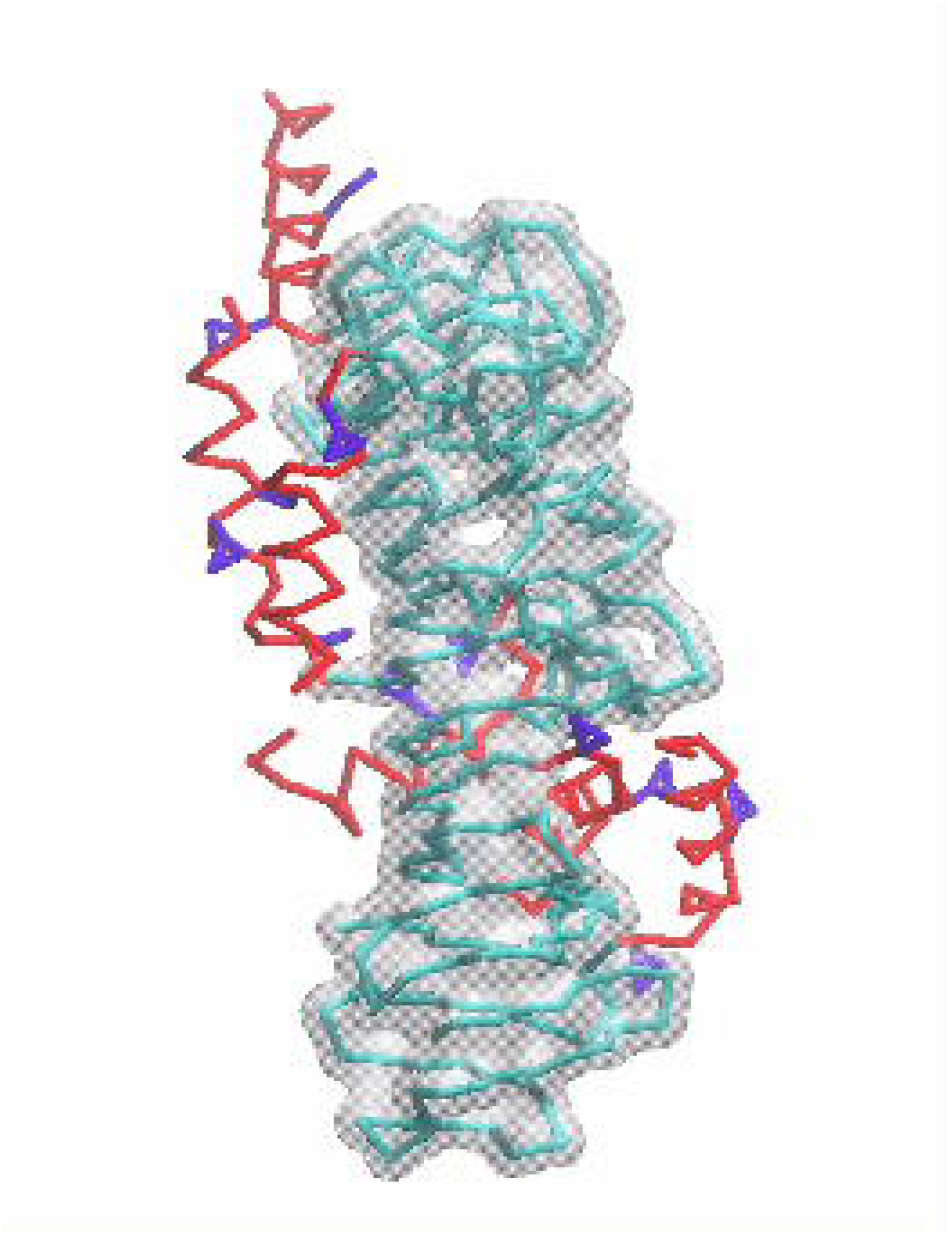
External distribution of N17 in the 45Q system. N17, polyQ and F are denoted in red, cyan and violet respectively.

## Acknowledgments

This research was supported by the National Science Foundation under grants CHE-1454948 and CHE-2202281. The authors acknowledge the University of Maryland supercomputing resources (http://hpcc.umd.edu) made available for conducting the research reported in this paper.

## Notes

### Competing Interest Statement

The authors have declared no competing interest.

### Summary of Updates

Title and methods section updated; Section on polymorphism observed in Huntingtin oligomers added to results.

