## Supplementary File for "Unraveling the Molecular Complexity of N-Terminus Huntingtin Oligomers: Insights into Polymorphic Structures"

### Supplementary Information for Unraveling the Molecular Complexity of N-Terminus Huntingtin Oligomers: Insights into Polymorphic Structures

Neha Nanajkar, Abhilash Sahoo, Silvina Matysiak

May 7, 2024

For the monomeric peptide systems, a single N17 + polyQ (polyQ= 7, 15, 35, 40, 45) was placed in a cubic water box of 8 nm length. These systems were minimized and equilibrated in the same manner as the aggregated peptide systems (see main text). Each simulation was run for 200ns with NVT ensemble. The first 50 nanosenconds were discarded as equilibration and the following 150 nanoseconds were reweighted to 380K for analysis.

Fig S1 presents the representative snapshots of each peptide system. The N17 domain is demarcated in red and the polyQ region in cyan. As the polyQ length increases, it is evident that a more collapsed conformation is assumed. In the 7Q and 15Q systems, the monomer favours more elongated conformations, while compact, collapsed conformations are seen with longer polyQ domains (polyQ=7,15,35,40,45). Fig S2 presents the potential of mean force with asphericity as a reaction coordinate. An asphericity value of 0 corresponds to a globular structure and a value of 1 indicates more planar conformations. The shift from elongated conformations towards globular conformations is apparent from the reduction in asphericity. This is in agreement with a number of studies that demonstrate the globular nature of the segments at longer polyQ lengths [1, 2].

Fig S3 presents backbone contact maps for the monomeric systems. In the N17 region, there is a high probability of i:i+4 contacts, indicative of helical structures. This persists in the polyQ region, although the probability of helix formation is comparatively lower. In Fig S3D, S3E we note the presence of contagious off-diagonal contacts in the polyQ domain, which is representative of intra-molecular beta-sheets. Although sheets were observed in the N17+35Q system, they were not as abundant. Figure S4 presents a closer look at the interactions of the polyQ domain. We notice an increase in the average number of Q-Q interactions per residue increases with increasing polyQ length. A glutamine residue in the N17+7Q system would interact with between 1-2 other glutamine residues. This increases to over 3 other glutamine residues for polyQ of 35,40 or 45. The increase in Q-Q connectivity can be associated with the preference of GLN residues to self-interact, forming collapsed structures that are linked through polar interactions(such as hydrogen bonding). Thus, we also see a corresponding decrease in the average number of Q-Water interactions. Taken together, we confirm that polyQ is a particularly flexible domain that is capable of assuming a variety of conformations, and assumes a more collapsed conformation at longer polyQ lengths .

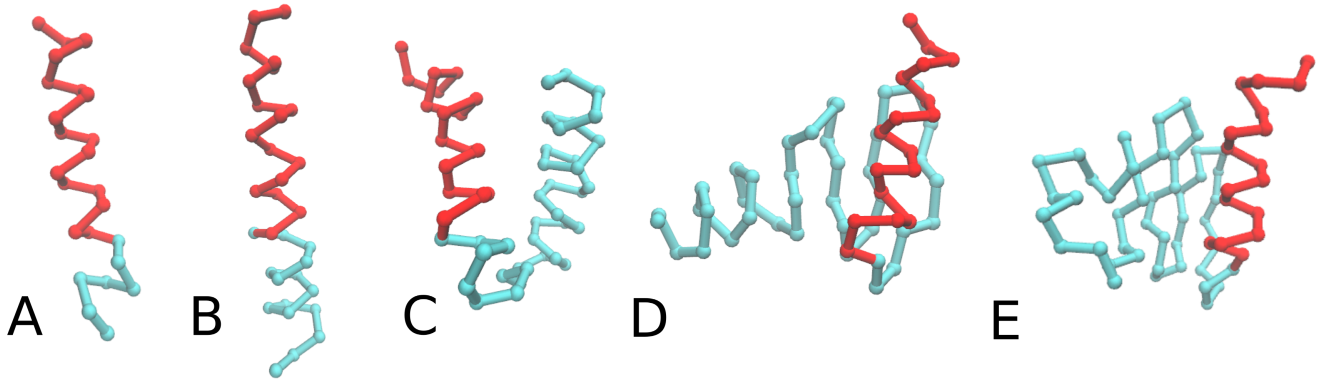


Figure 1: Representative snapshots of monomeric N17+ polyQ at 380K - A) N17 + 7Q, B) N17 + 15Q, C) N17 + 35Q, D) N17 + 40Q, E) N17 + 40Q. Snapshots selected are the corresponding asphericity values of the global minima, for each peptide system.

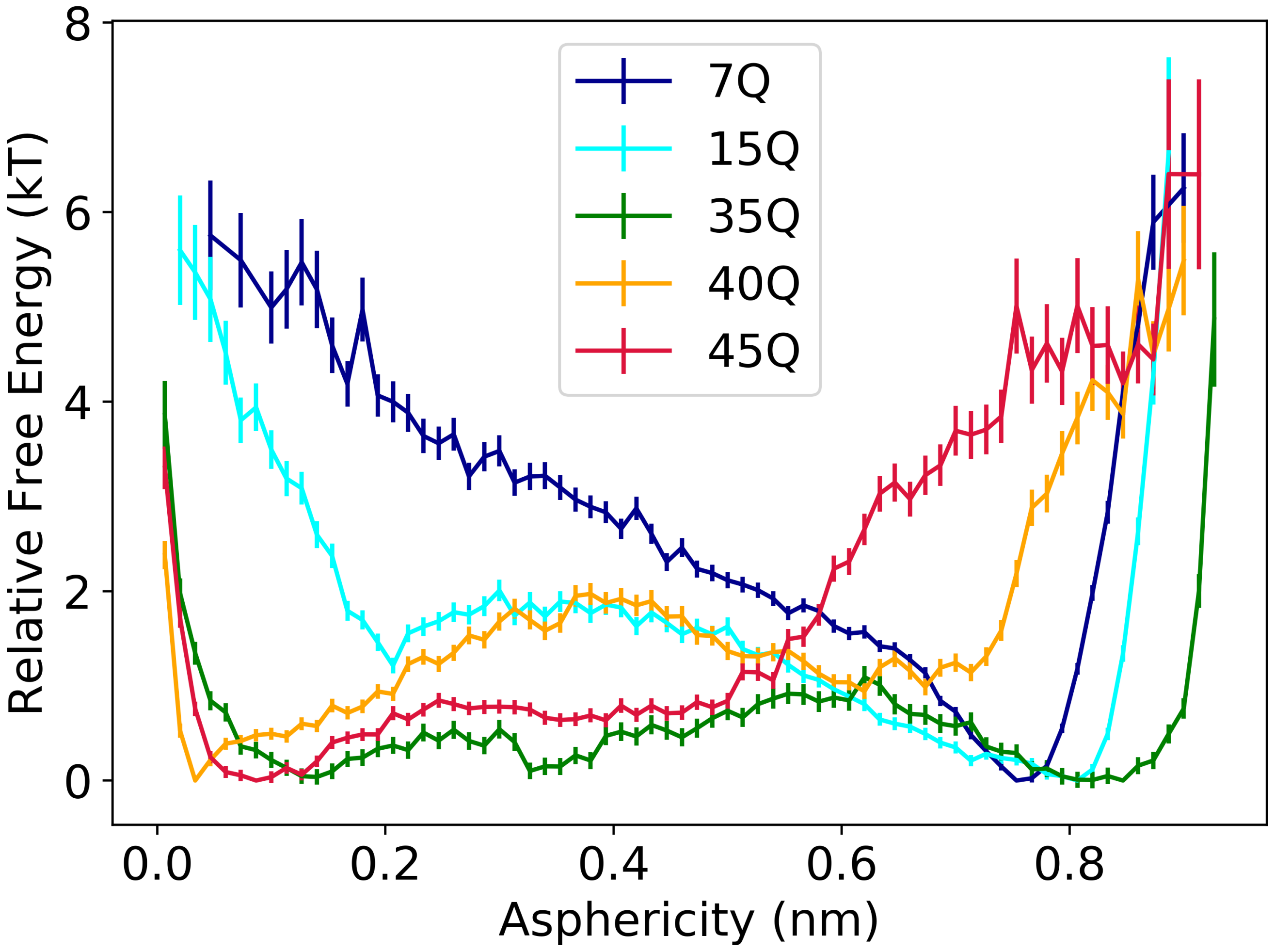


Figure 2: Potential of mean force of monomeric N17+ polyQ reweighted to T=380K, with asphericity as the reaction coordinate. An asphericity value of one indicates a more extended, planar structure, while a value of zero would indicate a a compact, spherical structure.

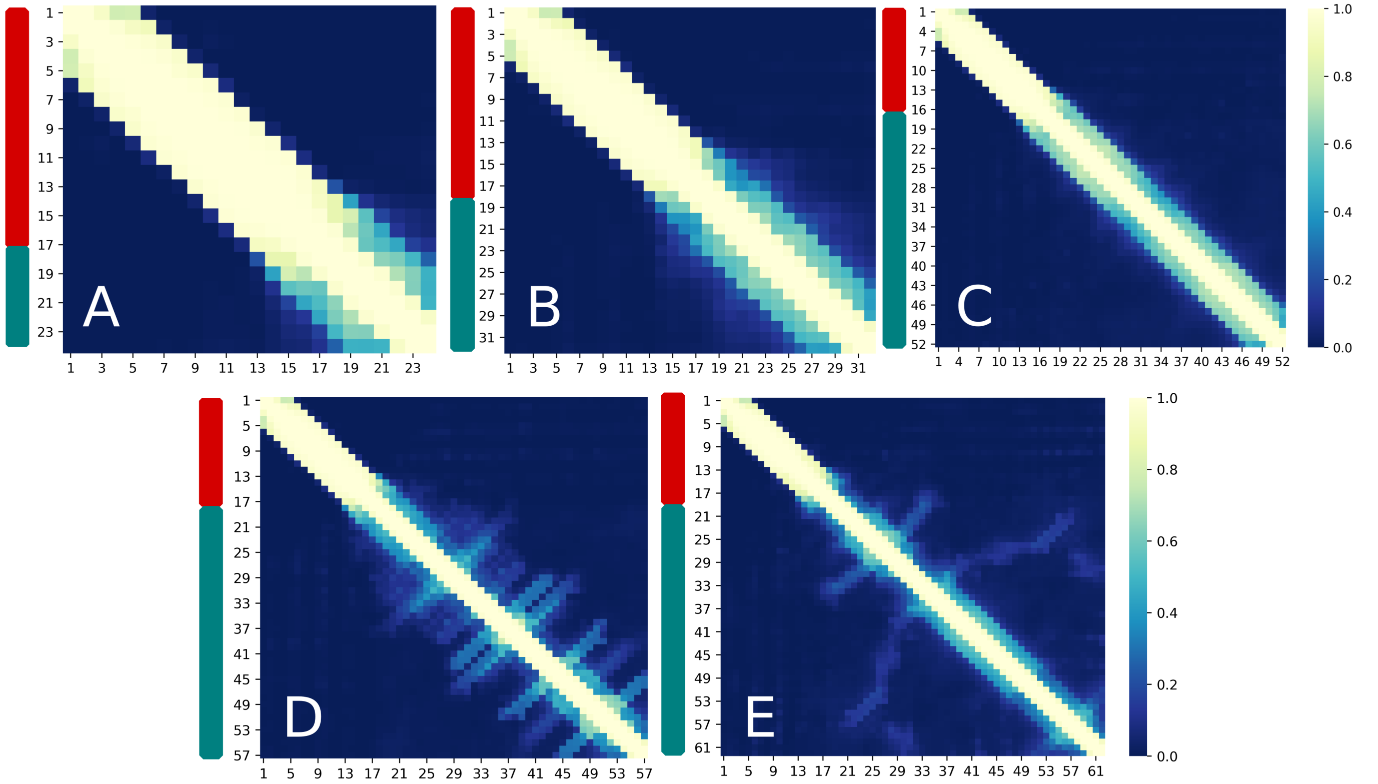


Figure 3: **Backbone contacts maps for single peptide systems -** A) N17 + 7Q, B) N17 + 15Q, C) N17 + 35Q, D) N17 + 40Q, E) N17 + 45Q, reweighted to T= 380K. Contacts are defined as backbone beads within 7 A° of each other. N17 and polyQ domains are marked in red and cyan respectively.

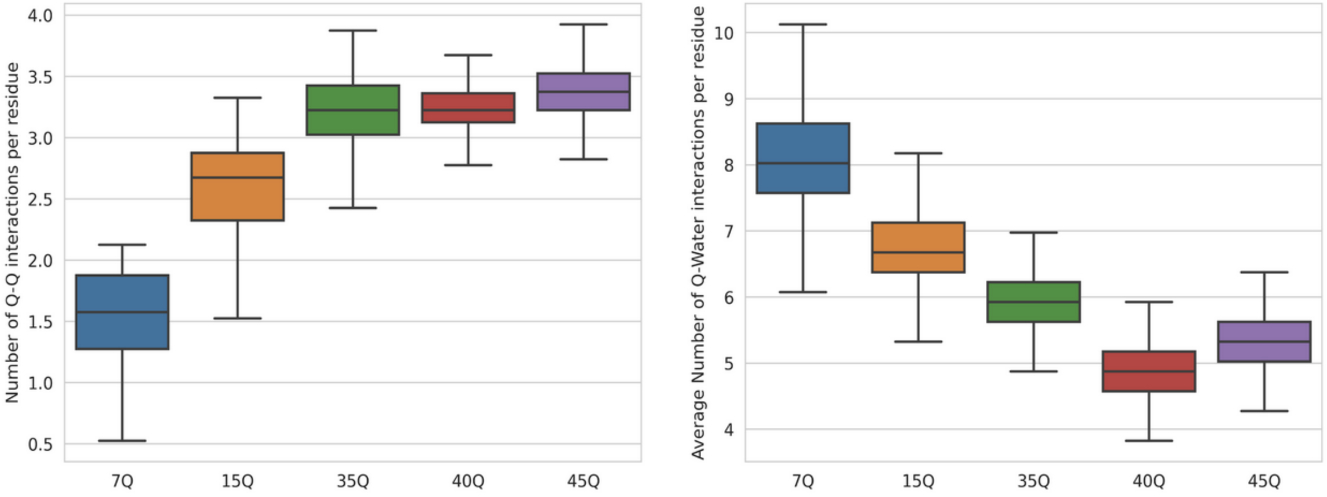


Figure 4: **Domain specific interactions of the polyQ domain in monomeric htt, reweighted to 380K.** A) Number of Q-Q interactions per residue for each peptide system. B) Number of waters interacting with each glutamine residue
